# Evaluating local assembly strategies for human long-read structural variant calling

**DOI:** 10.64898/2026.11.11.765824

**Authors:** Tim Mirus, Richard Lüpken, Birte Kehr

## Abstract

**Motivation:** Long-read sequencing enables detection of structural variants (SVs) with far greater accuracy and completeness than was previously possible. This advance has spurred the development of many SV callers that implement diverse algorithmic strategies. Commonly, SV calling proceeds in several processing steps. Evaluations of SV callers, however, typically focus on overall performance and conceal which steps drive observed differences. This hinders targeted, evidence-based improvements to existing SV calling approaches.

**Results:** Here, we implemented five consensus computation strategies similar to those employed by popular SV callers that perform a local assembly. We compare their outputs on extensive simulated and real data and reveal that some strategies are more effective for particular input data types. For example, the strategy employed by PacBio’s tool Sawfish works very well on HiFi reads and recent ONT data, but fails on the older and noisier ONT reads. While the read error profiles and the read pre-processing has a substantial influence on some strategies, other strategies are more robust overall. These findings guide algorithmic decisions for the development of improved future SV callers.

**Implementation and availability:** Source code is available at https://github.com/kehrlab/LR-Cons.

## Introduction

The last decade has seen substantial advances in our ability to detect genomic structural variants (SVs) [Logsdon et al., 2025]. Much of this progress has been driven by long-read sequencing technologies [Chaisson et al., 2015]. Despite the ability of long reads to entirely span relatively large variants and reveal previously invisible variants, reliably detecting and precisely characterizing the full spectrum of SVs in routine genome analysis continues to pose a challenge [Collins and Talkowski, 2025]. Developing improved methods to detect and characterize SVs in long-read data remains an active area of research [Ahsan et al., 2023, Olson et al., 2023].

Broadly, existing germline SV callers for long-read sequencing data aligned to either a linear or a pangenome reference genome can be grouped into different classes: Clustering-based methods detect SVs by collecting and aggregating SV signatures in read alignments [Heller and Vingron, 2019, Jiang et al., 2020, Smolka et al., 2024]; local assembly-based approaches additionally assemble reads within candidate loci to produce consensus sequences of local haplotypes that are then aligned to the reference to resolve and characterize SVs [Denti et al., 2023, Chen et al., 2023, Saunders et al., 2025, Jiang et al., 2026]; whole-genome *de novo* assembly-based strategies reconstruct (often haplotype-aware) assemblies of the sequenced whole genome and compare them to the reference [Li et al., 2018, Heller and Vingron, 2020, Ebert et al., 2021]; finally, deep-learning-based tools encode SV signatures observed in reads in a matrix or image that allows prediction of SVs through deep learning models [Lin et al., 2022, Ding and Luo, 2022, Popic et al., 2023, Wang et al., 2025].

Local assembly-based approaches currently provide a favourable trade-off between resource demands, including data and compute, and the accuracy of SV calling. Indeed, PacBio recently replaced their clustering-based method PBSV with the local-assembly approach Sawfish [Saunders et al., 2025] in its recommended variant analysis pipeline (https://github.com/PacificBiosciences/HiFi-human-WGS-WDL). These approaches are therefore currently particularly likely to be adopted in practice (e.g. in clinical genome analysis).

While the available callers using local assembly differ markedly in methodology, all of them broadly follow the same sequential steps for SV calling from aligned long-read data: (i) identification of candidate SV locations and SV-supporting reads, (ii) consensus computation at candidate loci, and (iii) SV detection from consensus sequences by comparison to the reference genome. As evaluation and benchmarking of SV callers is generally conducted on the level of final SV calls, it remains unclear how implementation details at each step affect overall performance, and whether differences in results are driven more by the broad approach or by choices made within individual steps. To develop improved methods in a well-founded way, we need an understanding of how the choice of algorithm for each individual step impacts SV calling performance.

For consensus computation at candidate SV loci (step (ii)), most local assembly approaches use partial order alignment (POA) [Lee et al., 2002]. POA extends the classic dynamic programming approach for pairwise sequence alignment [Needleman and Wunsch, 1970] by building a partial order graph representing a multiple sequence alignment. Using the heaviest bundle algorithm [Lee, 2003], we can then retrieve one or more consensus sequences from the POA graph. However, when studying how POA is integrated into existing SV callers, it becomes apparent that the developers had to make fundamental algorithmic choices apart from selecting default parameter values. In its simplest form, POA takes as input a single set of sequences and computes a single consensus sequence. In contrast, the reads aligning to most loci in the human genome originate from two copies of the chromosome, the two haplotypes. A key challenge for consensus computation in SV callers is to separate the haplotypes. This can be carried out before, during, or after POA, and for each case multiple implementation strategies are possible. For example, Sawfish computes the POA using a variable number of partial order graphs [Saunders et al., 2025], separating the haplotypes during POA. DeBreak first assigns SV-supporting reads to one or two clusters based on estimated SV size, and then computes a single POA per cluster [Chen et al., 2023]. Similarly, SVDSS computes two POAs after clustering, but uses a sequence similarity score for clustering read superstrings [Denti et al., 2023]. It remains unclear why the precise approaches were chosen and how informed the decisions for the particular POA strategy were.

In this work, we set out to compare and evaluate strategies for building consensus sequences similar to those implemented in prominent local assembly-based SV callers. We (re-)implemented a total of five POA strategies for local haplotype assembly at SV candidate sites and evaluate them on extensive simulated and real data. Although we are zooming in on only one step in the SV calling workflow, we identify significant differences in robustness to input data quality for reconstructing accurate consensus sequences, which is the basis for subsequent variant characterization. Our results emphasize the importance of informed algorithmic choices at each individual step of complex computational tasks such as SV calling.

## Methods

Our evaluation of distinct strategies for consensus computation at SV loci in the human genome is based on simulated and real long-read data (see *Preparation of input data*). We provide identical sequence sets per variant locus as input to all strategies (see *Extraction of variant-specific sequences*). We compare five distinct consensus-generation strategies (Figure 1) and, for simulated data, a baseline strategy that has knowledge of the true read phase (see *Consensus sequence generation strategies*). We evaluate the success rate of consensus sequence reconstruction at homozygous and heterozygous SV loci (see *Performance Evaluation*).

**Fig 1.**
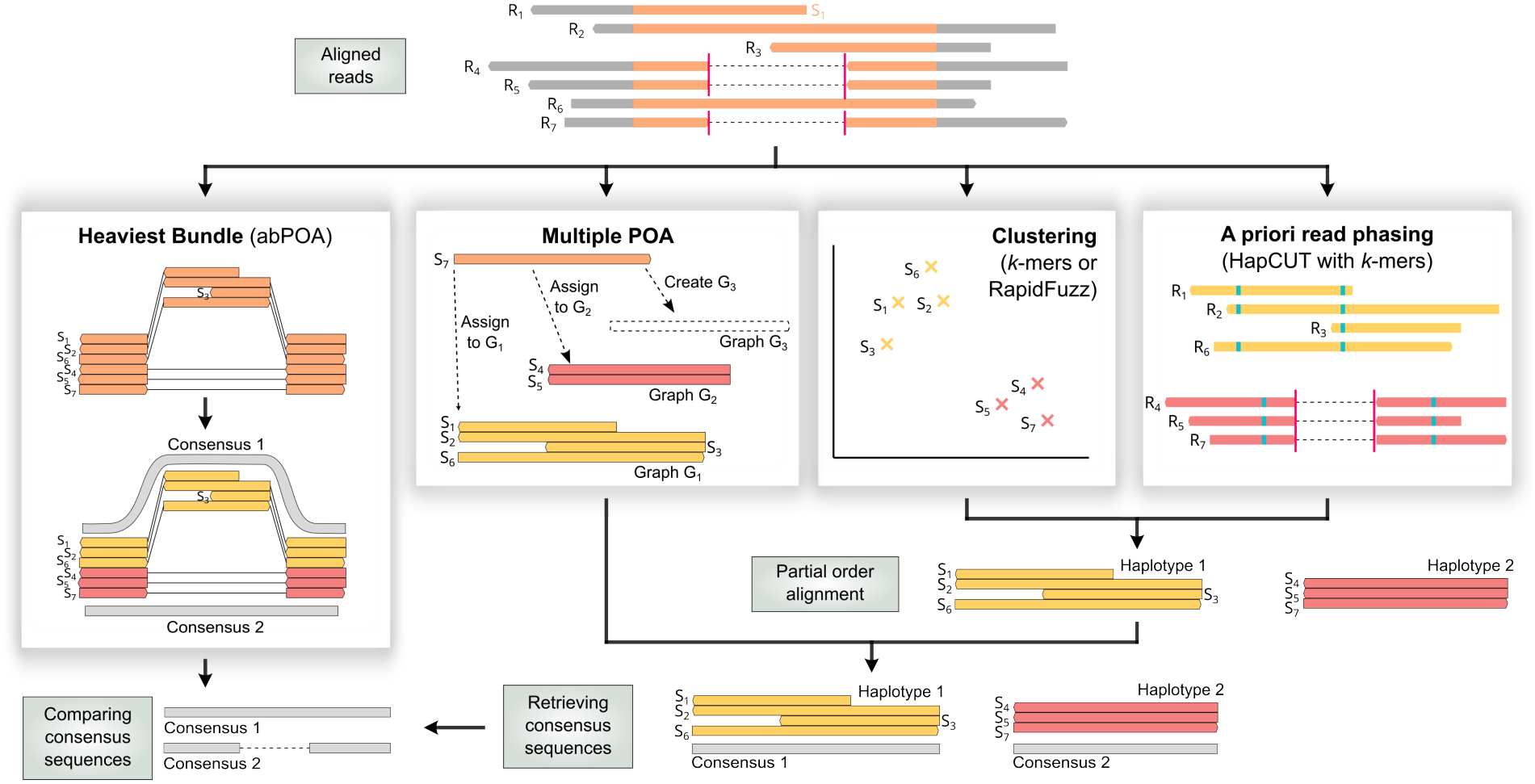
Overview of the five implemented consensus sequence generation strategies. *R*_*i*_ represent reads for a given variant locus and *S*_*i*_ are extracted subsequences. Yellow and red indicate the haplotype assignment of a sequence.

### Preparation of input data

We use PacBio HiFi and ONT long read sequencing data from HG002 in conjunction with the GIAB v5.0q callset for our evaluation (see *Data Availability*). For both technologies, we downloaded the latest available data sets (`HiFi 2025, ONT` 2025), as well as older data (`HiFi 2020, ONT` 2020). HiFi 2025 has the lowest sequencing depth (32.51*×*). We therefore used samtools [Danecek et al., 2021] to subsample the other data sets to a coverage of approximately 32.5*×*, and additionally subsampled all data sets to 15*×* and 20*×*. To obtain a high-confident callset to be used as ground truth for HG002 SVs we filtered the GIAB callset by excluding regions containing more than one SV and removing variants smaller than 50 bp. We also removed SVs above 5 kbp in order to focus on variants that can be expected to be spanned by entire reads in all used data sets.

In addition, we reconstructed HG002 haplotypes in-silico by inserting all properly phased SNPs and the remaining SVs into the T2T-CHM13 reference genome and simulated long reads using PBSIM3 [Ono et al., 2022], which has been benchmarked favourably against other tools for ONT data simulation [Taouk et al., 2026]. We used quality score models with default parameters for ONT and HiFi read generation, respectively, to generate sequencing data at 10*×* and 15*×* coverage per haplotype. Target sequencing accuracy was set to 99% for ONT reads, while PacBio data were generated with ten passes and then further processed with ccs v6.4.0 to obtain HiFi reads.

Finally, we simulated an artificial diploid chromosome with random, uniformly distributed SVs. We introduced a total of 16000 SV loci in non-telomere and non-centromere regions of two chromosome 1 (chr1) copies from the T2T-CHM13 reference genome, and randomly substituted single base positions to simulate SNVs (see Supplementary Note 1). Reads from the two haplotypes were simulated with PBSIM3 as described above, with an additional coverage level of 30*×* per haplotype.

We aligned both the simulated and real long reads to the T2T-CHM13 reference genome using minimap2 [Li, 2018, 2021] with the parameter sets for ONT (`-x map-ont`) and HiFi (`-x map-hifi`) reads. In order to reduce differences between datasets caused by random coverage fluctuations, we identified GIAB SV regions that have low coverage (*<* 6*×*) or an insufficient number of reads with SV signals (*<* 3) in any of the four real data sets at 32.5*×* coverage and removed the corresponding SVs from our evaluation. The remaining set of loci contains 13506 SV regions (6873 deletions and 6633 insertions).

### Extraction of variant-specific sequences

We make the assumption that all variant loci *v* can be detected in an initial step of the SV calling workflow, including either perfect estimates of their start and end positions, *x*_*v*_ and *y*_*v*_, in the reference genome, or imperfect estimates where we add Gaussian noise (i.e. 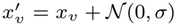 and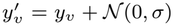), where *σ ∈ {*0, 50, 200*}*. We extract locus-specific subsequences *S*_*i*_ from read alignments *R*_*i*_ to avoid assembling irrelevant sequence segments and enforce similarity of sequence ends for easier alignment. The resulting sequence sets are supplied as input to every consensus-generation strategy described below.

Defining *f* as the length of a flanking region, we collect the set of reads *ℛ* that overlap the reference interval *I*_*v*_ = [*x*_*v*_ *− f, y*_*v*_ + *f*]. Unless otherwise stated, we use *f* = 500 bp. To ensure consistent read orientation, we query the orientation of read alignments at the the boundaries of *I*_*v*_ and reverse complement reads if necessary. Next, we extract a subsequence *S*_*i*_ = (*R*_*i*_, *p*_*i*_, *l*_*i*_) from each read *R*_*i*_ *∈ ℛ* where *p*_*i*_ is *S*_*i*_’s start position in *R*_*i*_ and *l*_*i*_ is *S*_*i*_’s length. We traverse the CIGAR strings of *R*_*i*_’s alignments to define *S*_*i*_ as the smallest subsequence of *R*_*i*_ such that all positions of *R*_*i*_ that align within *I*_*v*_ are contained in *S*_*i*_. We define the set *S* as the subsequences *S*_*i*_ of reads *R*_*i*_ that are anchored in the flanking region on both sides of the variant. Furthermore, we define *S*^+^ as an extended set (*S ⊆ S*^+^) that may contain subsequences of reads anchored only on one side of the variant. Given a minimum length *ℓ* of the sequences in *S*, we crop the reads that are anchored only at one side at most *ℓ* bp away from the anchor. For subsequent processing steps, we assume that the sequences in *S* and *S*^+^ are ordered from longest to shortest such that *l*_*i*_ *≥ l*_*i*+1_, to ensure robust performance when iteratively adding the *S*_*i*_ to a POA.

### Consensus sequence generation strategies

Figure 1 provides an overview of the five evaluated strategies for consensus sequence generation. If not otherwise stated, we implemented POA using the SPOA library [Vaser et al., 2017], which provides an established and reliable implementation of the original POA algorithm. We evaluate local and semi-global alignment of the sequences *S*_*i*_ *∈ S*^+^ as well as global and local alignment of the sequences *S*_*i*_ *∈ S* against the POA graph. Across all strategies, we derive consensus sequences from a POA only if it was constructed from at least three input sequences. We discard any consensus sequence that is much shorter than expected (smaller than *f* bp). After initially testing several different alignment scoring schemes (Supplementary Note 10), we settled on a unit-cost scoring scheme with linear gap penalties (match = 1, mismatch = *−*1, gap = *−*1), which showed consistent results across strategies and alignment modes.

All strategies output up to two consensus sequences per locus, but some strategies enforce exactly two consensus sequences. To ensure a fair evaluation of the ability to reconstruct distinct SV alleles, we align each strategy’s proposed haplotypes to each other (*Comparing consensus sequences* in Figure 1). If the sequence similarity is above a pre-defined threshold, we only keep the longer consensus sequence. We calculate the sequence similarity as 1 *− d* with *d* being the Levenshtein distance divided by the shorter consensus sequence’s length (*l*_*s*_) and define the threshold as 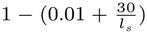, allowing for 1% divergence and up to 30 bp difference due to smaller indels between sequences representing the same SV. We implemented this comparison using semi-global alignment from the SeqAn C++ library v2.4 [Reinert et al., 2017] with a Levenshtein distance scoring scheme.

#### Heaviest bundle algorithm using abPOA

The first strategy in our evaluation creates a single POA graph from the sequences *S* or *S*^+^ of one candidate variant locus and subsequently retrieves up to two consensus sequences from the graph, to represent two potentially distinct structural variant alleles. Since the SPOA library allows retrieval of only a single consensus sequence, we use abPOA [Gao et al., 2021] with parameters *w* = 6, *k* = 9, and *min w* = 10 to construct a POA graph and retrieve consensus sequences. We enable abPOA’s minimizer-based progressive tree feature to increase performance. We note that the implementation of the heaviest bundle algorithm in abPOA [Gao et al., 2021] deviates from the original publication [Lee, 2003] in a way that avoids phase switch errors. abPOA detects and clusters sequences based on heterozygous bases in the POA graph, before extracting cluster-specific consensus sequences from the graph with a variation of the heaviest bundle algorithm.

#### Sawfish’s multiple POA graph strategy

The SV caller Sawfish [Saunders et al., 2025] assigns sequences to haplotypes during POA by building multiple POA graphs. When adding a sequence *S*_*i*_ to the alignment of *S*_1_, …, *S*_*i−*1_, Sawfish decides based on alignment score whether to add *S*_*i*_ to one of the existing POA graphs or initialize a new graph. By default, Sawfish builds at most eight POA graphs. Eventually, Sawfish retrieves consensus sequences only from the two graphs with the highest supporting read count.

We implemented the same basic strategy for our evaluation. Similar to Sawfish, we assign a sequence from *S* or *S*^+^ to an existing graph if more than 95% of the sequence is locally aligned and its alignment score normalized by alignment length exceeds 0.96. If multiple graphs fulfil both alignment quality thresholds, we align a sequence to the graph with the higher alignment score. If no graph meets the alignment requirements for a given sequence, we initialize a new graph or, if the maximum number of 8 graphs is reached, discard the sequence.

#### Similarity-based sequence clustering

A straightforward strategy for generating multiple consensus sequences is to first cluster input sequences before building a POA graph per cluster and retrieving one consensus sequence per POA graph [Chen et al., 2023, Denti et al., 2023]. In our evaluation, we included a simple alignment-free strategy that uses a *k*-mer-count spectrum for clustering and an alignment-based strategy that uses a sequence similarity score for clustering.

Our alignment-free strategy clusters sequences in *S* or *S*^+^ based on *k*-mer counts, with *k* = 11 for ONT and *k* = 21 for HiFi data (Supplementary Note 6). Without loss of generality, we use sequence set *S* in the following paragraph. We exclude each *k*-mer *a* that has a count *n*_*ia*_ *>* 100 in any sequence 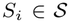, and each *k*-mer *b* with an insufficient total count 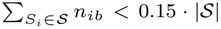. Subsequently, we select the 100 *k*-mers with the highest count variability across all sequences for k-means clustering with two clusters.

Our alignment-based strategy was inspired by SVDSS [Denti et al., 2023], which clusters its superstrings based on sequence size and similarity, with the similarity being calculated using RapidFuzz (https://github.com/rapidfuzz/RapidFuzz). RapidFuzz defines the similarity Sim_*L*_(*S*_*i*_, *S*_*j*_) for two sequences *S*_*i*_, *S*_*j*_ *∈ S* via the Levenshtein distance. As input sequences are not always of similar length, we define

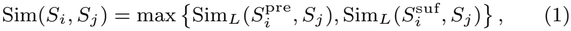

where *S*_*j*_ is the shorter sequence and 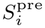 and 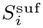 refer to the prefix and suffix of length |*S*_*j*_ |, respectively. To avoid the computational cost for calculating all-against-all pairwise similarities, we heuristically determine one representative sequence per haplotype and assign all other sequences to the most similar haplotype representative. In particular, we choose the longest sequence 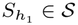 as representative of the first haplotype. To select the second haplotype representative, we compute the similarities Sim 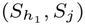 of all 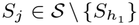 and choose 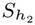 as the sequence at the first quartile of the resulting similarity distribution. We further compute the similarity of all 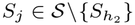 to 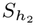. Finally, we assign each sequence *S*_*i*_ *∈ S* to the more similar representative if *S*_*i*_ is at least as similar as the first quartile of the similarity distribution to either 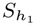 or 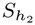.

After clustering with either strategy, one POA is computed per cluster and one consensus sequence retrieved per POA.

#### A priori read phasing

All strategies described above distinguish between the two alleles at SV loci solely based on the similarity of the read subsequences *S*_*i*_ *∈ S* or *S*_*i*_ *∈ S*^+^. Our last POA strategy locally phases the full read sequences *ℛ* in intervals of 100 kbp around SV loci, leverages the phase information to partition the *S*_*i*_, and computes a POA and consensus sequence for each partition. We implemented the idea of the HapCUT [Bansal and Bafna, 2008] phasing algorithm but, in contrast to HapCUT, determine the read phase based on “heterozygous” *k*-mers instead of input small variant calls.

More precisely, we count *k*-mer occurrences (*k* = 55 for ONT, *k* = 77 for PacBio) across all reads that align within the 100 kbp interval. We say that two different *k*-mers are *neighbours* if they have a Hamming distance of 1. For every *k*-mer Ref that occurs within the 100 kbp interval in the reference genome, we compare its count *c*^Ref^ to the counts *c*^Alt^ of all its neighbours Alt. A *k*-mer pair (Ref, Alt) with (*c*^Ref^, *c*^Alt^) is used for phasing if Alt is the only neighbour of Ref that satisfies max {*c*^Ref^, *c*^Alt^ */*} min {*c*^Ref^, *c*^Alt^} *<* 1.5 (i.e. the two counts differ by less than 1.5-fold).

Similar to HapCUT, we then partition the Ref of all selected *k*-mer pairs into two haplotypes by optimizing a global phasing score (Supplementary Note 4). Finally, we assign the input reads to haplotypes based on the majority vote of *k*-mers they contain.

#### Ground truth phasing

In our evaluation on simulated data below we included an additional baseline strategy for comparison, which uses the known phase of our simulated reads to partition the subsequences *S*_*i*_ in two groups, computes a POA per haplotype and derives two consensus sequences from these POAs.

### Performance Evaluation

We evaluate the output of each strategy at a given variant locus by aligning the created consensus sequences against the two ground-truth haplotype sequences at the SV locus (covering the entire variant and 2000 bp up- and downstream) using semi-global alignment with a Levenshtein distance scoring scheme (match = 0, mismatch = *−*1, gap = *−*1). For simulated data, the ground-truth sequences are known and extracted during haplotype creation, while for the real data, we reconstruct these sequences from the SV sequences given in the VCF file and flanking sequences extracted from the reference genome. Each consensus sequence is assigned to the haplotype with the higher-scoring alignment. We discard consensus sequences that do not align across a breakpoint position on the ground truth sequence or have a match rate *<* 95%.

#### Precision and recall

Based on the zygosity of the ground-truth SV, the number of consensus sequences generated, and the assigned haplotypes, we categorise the output into combinations of true positives, false positives and false negatives as shown in Figure 2a. We then use the categories to assess a strategy’s performance in terms of how many correct SV alleles can be reconstructed (recall) and how many erroneous or additional sequences are generated (precision). For each strategy, we count the size |*X*| of each category *X* and calculate for assessing heterozygous variants

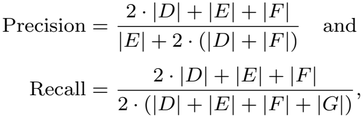

and analogously for homozyogous variants (Supplementary Note 3).

**Fig 2.**
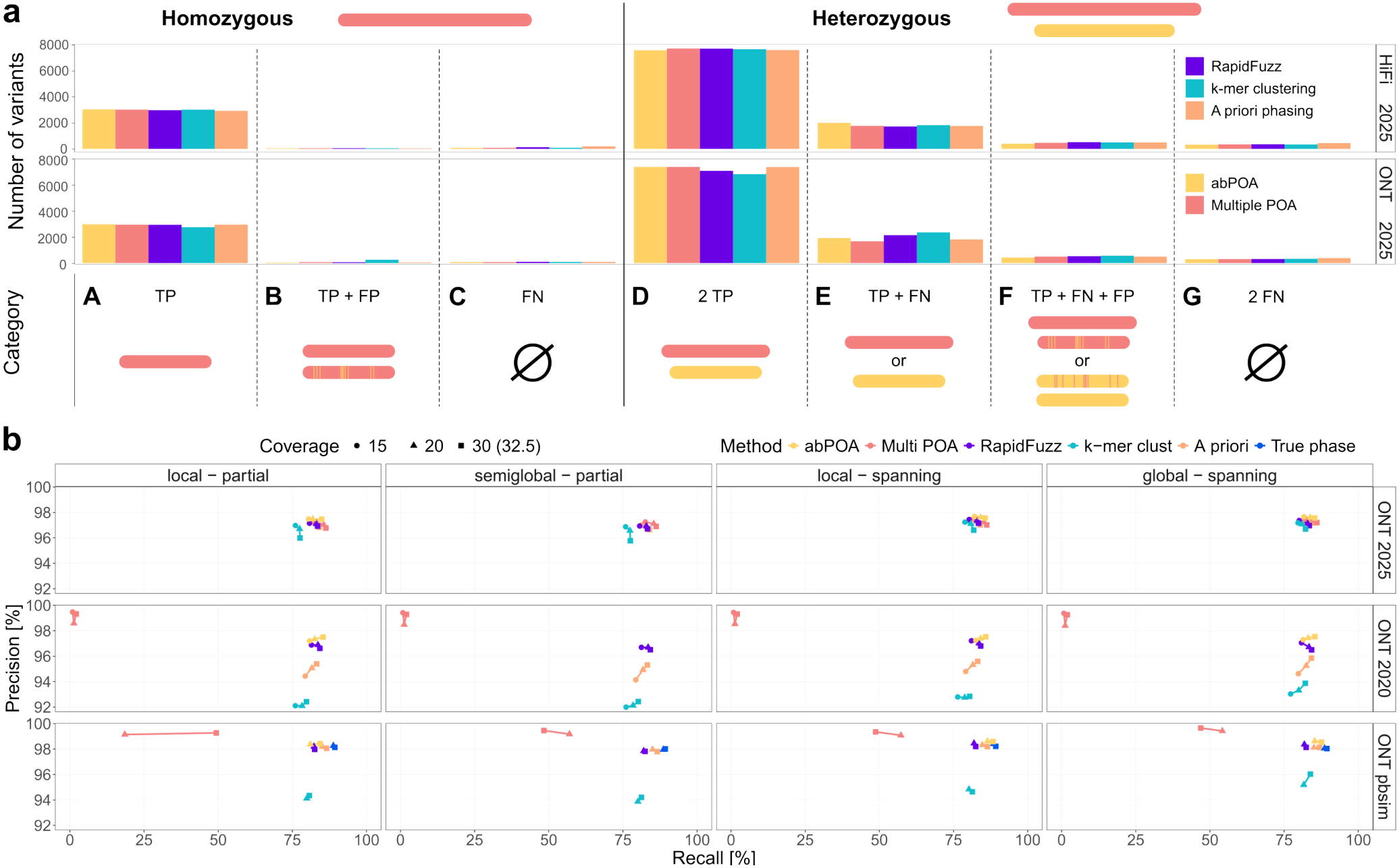
**(a)** Number of variants per category for real data at 32.5*×* coverage with the input read set *S* and global alignment. Top row indicates the true genotype of the simulated variant and bottom row indicates the configuration of constructed consensus sequences within a category. *TP*, True positive; *FN*, False negative; *FP*, False positive. **(b)** Precision-recall curves across coverages for all strategies at heterozygous GIAB variant loci. Rows show different ONT long read data sets (real: 2020, 2025; simulated). Columns show different combinations of input sets (spanning: *S*, partial: *S*^+^) and alignment mode. Precision and recall were calculated as described in *Performance Evaluation*.

## Results

We evaluate the different consensus computation strategies in various different scenarios defined by input data type and preprocessing, as well as alignment type. In all tested combinations, we calculated recall and precision based on the seven categories defined above.

### All strategies perform well under optimal conditions

Figure 2a displays the number of variants per strategy that fall into each of our seven categories when focussing on a single scenario that uses 32.5*×* coverage of the `ONT 2025` and `HiFi` 2025 data sets with global POA, and excluding partially overlapping reads to minimize variability in the input data. Overall, we observe few variant loci yielding no consensus sequence (categories *C* and *G*).

For homozygous variant loci (*n* = 3183), our implementation of the abPOA, multiple POA, RapidFuzz clustering, and a priori read phasing strategies show excellent performance in identifying homozygous loci and returning only a single consensus sequence (recall *>* 93%, precision *>* 97%). Only our *k*-mer-based clustering leads to a considerable amount of incorrect additional sequences on ONT data (|*B*| = 266, precision 92%).

For heterozygous loci (*n* = 10323), we observe slightly more differences between the strategies and sequencing technologies. On PacBio HiFi data all methods show very good performance with the best (multiple POA: |*D*| = 7722) and worst (abPOA: |*D*| = 7577) performing strategies still being close together. The performance on ONT data is more varied: Multiple POA (|*D*| = 7767) and abPOA (|*D*| = 7602) show a slight increase in performance compared to PacBio data, while RapidFuzz (|*D*| = 7248) and especially the *k*-mer based clustering strategy (|*D*| = 6993) resolve fewer loci correctly. All methods show a lower recall on heterozygous loci (82.2% *−* 86.1%) than on homozygous loci (96.8% *−* 97.5%).

These results indicate that variation in input data and setup may influence performance in different ways, while the peak performance of all strategies on high-quality input data is comparable.

### Effects of data quality, input sequence set, and coverage

Placing the focus on the strategies’ ability to correctly separate two distinct alleles, we restrict all following analyses to the more challenging heterozygous variant loci. Figure 2b shows the results on the GIAB SVs for the different ONT datasets when running local, semi-global, or global POA inside all strategies using only the set *S* (spanning) or the set *S*^+^ (partial), summarizing the seven categories in precision and recall values. Global POA requires input sequences to have similar lengths, so we only tested global POA when restricting the input to those reads that fully overlap the variant locus (spanning). Furthermore, abPOA currently does not support semi-global alignment. Performance on PacBio HiFi data remains largely stable across data sets (Supplementary Figure 11).

When using input set *S*, performance on the most recent `ONT 2025` dataset is on par with PacBio HiFi data as also shown in Figure 2a. At 32.5*×* coverage, recall values range between 82% (k-mer clustering) and 86% (Multiple POA) and precision values from 97% (k-mer clustering) to 98% (abPOA). When switching to the *S*^+^ input sets, we note a drop in recall for the k-mer based clustering, while the other strategies remain unaffected.

However, this performance does not translate to older data. While abPOA and RapidFuzz show almost unchanged performance on the `ONT 2020` data, the k-mer based clustering and a priori phasing – which also relies on k-mers – both suffer due to the reduced accuracy of the older ONT data, with precision and recall dropping to 95% and 83% for the phasing strategy and even further to 92% and 80% for the k-mer based clustering at 32.5*×* coverage. The most notable difference can be observed for multiple POA, which showed the highest recall on the the ONT 2025, dropping to 2% on the older dataset. We suspect that the higher error rate causes the iterative alignment of reads to POA graphs with merging threshold 96% to fail. Repeating the experiment with a lowered merging threshold of 90% confirms this (Supplementary Figure 13), but the lowered threshold fails to separate distinct alleles at heterozygous loci, causing the performance to stay well below competing methods on this data set.

The simulated ONT data – PBSIM’s quality score model was trained on older ONT data – confirms our observations for the multiple POA and k-mer clustering strategies. The performance drops are less pronounced, indicating higher data quality for the simulated data to an extent that allows the a priori phasing to rank among the best methods again.

The difference between local and semi-global alignment for the partial input set and between global and local alignment for the spanning input set are minor across all strategies. As expected, higher sequencing depth generally leads to higher recall. With global alignment and input sequence set *S*, recall on the `ONT 2025` or `HiFi 2025` data increases by 2.5% (*k*-mer clustering) to 6.5% (RapidFuzz) when increasing coverage from 15*×* to 30*×*.

The length of flanking sequences extracted around potential loci is another variable with a potential impact on consensus sequence generation. SV signals in read alignment, especially in low-complexity regions, can be scattered around the actual breakpoint (Supplementary Note 8). Increasing the length of sequences extracted from reads for consensus sequence construction (*f* = 1000 bp) increases recall for all strategies except multiple POA, as shown in Figure 3a. This is likely due to larger flanking regions increasing the chances of capturing local variation completely. Interestingly, larger flanking regions have an overall detrimental effect on recall on our simulated dataset. This effect decreases with increased breakpoint uncertainty (Figure 3b).

**Fig 3.**
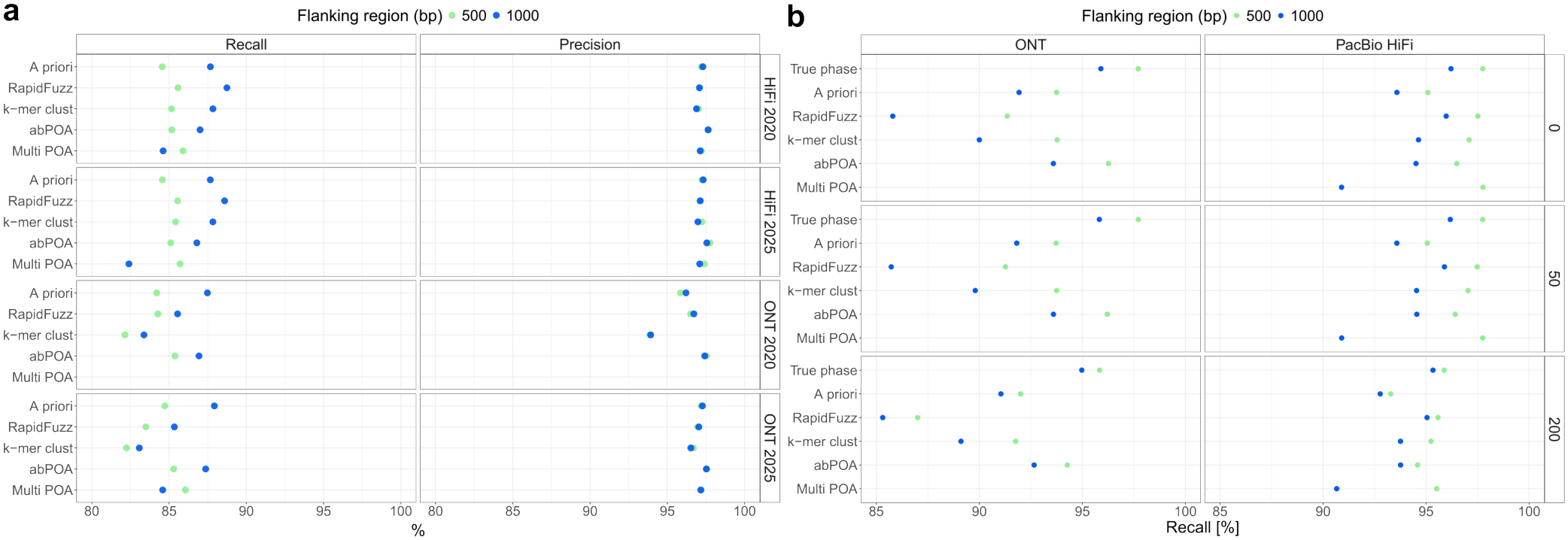
Impact of flanking region size on consensus sequences (input sets *S*, global alignment). **(a)** Precision and recall on real data. Multi POA on `ONT 2020` data is out of scope and therefore excluded. **(b)** Recall on simulated data, depending on technology and level of Gaussian noise (*σ*) added to breakpoint positions (rows).

## Discussion and conclusion

In this work, we studied different strategies for computing consensus sequences at SV loci from long reads using POA and assessed the impact of several key variables on their performance in the long read SV calling problem. We found that data quality and preparation of input data are the dominant factors, influencing our strategies’ performances in different ways, while sequencing depth has a consistent influence on recall across almost all scenarios and strategies.

On the most recent ONT and PacBio HiFi data sets, all tested strategies performed on the same high level, highlighting the substantial improvements in data quality ONT long reads have made in recent years, and suggesting that with sufficiently high data quality the exact choice of algorithm for consensus sequence generation becomes secondary. On older data, our comparison shows marked differences for some strategies in behaviour between the two sequencing technologies. The only strategies that consistently perform well across all evaluated setups are abPOA and RapidFuzz, with abPOA generally achieving slightly higher recall and precision. Performance of all strategies on older HiFi data sets is on par with the more recent data, while on older ONT data sets performance varies greatly, with k-mer based approaches suffering noticeable drops in precision. The recall of the multiple POA strategy, which narrowly outperformed other strategies on recent ONT data, drops to almost zero because the sequence-to-graph assignment breaks down at higher error rates. Lowering the assignment threshold leads to subpar performance, too, indicating that extensive, potentially dataset specific, finetuning of parameters may be needed to make this approach effective for lower-quality input data. Overall, we observe more stable results with slightly higher precision when using only those reads as input that span the entire variant and can be anchored to the reference on either side.

We investigated the effect of increasing the length of the flanking regions included in the assembled sequences and observe a clear positive effect on the recall of all strategies except multiple POA on real data. We assume this to be due to a combination of shifted breakpoint positions in the GIAB callset compared with our long read alignment data and high variability of breakpoint positions observed for some SVs, especially in repetitive genomic regions. This hypothesis is supported by the fact that we see a slight detrimental effect on recall when increasing flanking region size from 500 bp to 1000 bp on simulated data without breakpoint uncertainty and without SVs in centromere or telomere regions. This effect decreases when adding uncertainty to the known breakpoint positions in the simulated data. We attribute this to longer input sequences introducing noise into the generated consensus sequences, leading to some unmatched sequences. The latter effect is likely obscured by the uncertainties contained in real data sets.

The example of flanking sequence length illustrates the difficulties inherent in evaluating tools and methods using existing SV callsets as ground truth. Even after extensive filtering of the GIAB v5.0q callset, we have no guarantee as to the correctness of the SV positions and consequently the allele sequences we align our generated consensus sequences against. Differences in SV representation between long read alignments and alignments of whole genome assemblies may also influence results on real data. Our simulated data can mitigate this issue by providing data sets with accurate ground truth sequences for evaluation.

Our results also reflect the inherent limitations of SV calling based on local assembly, as recall remains below 86% even on the filtered set of SV loci, with up to 60% of false negative calls falling into centromere regions (Supplementary Figure 6). We do not evaluate the impact of the different POA-based local assembly strategies on the final SV calls here, but it is clear that a missing and inaccurate consensus sequence cannot result in an accurate SV call.

Despite our aim to highlight the importance of well-founded decisions during algorithm development, our implemented strategies and evaluation setup are not free of ad hoc decisions and parameters that have not been investigated to their fullest extent. Among these are the accuracy of the simulated data, the limited investigation of POA scoring schemes, and thresholds for sequence merging and required read support per haplotype. Similarly, none of the implemented strategies was fully optimized. Instead, we kept thresholds and key parameters comparable to isolate the effects of major variables on strategy performance. Further parameter optimizations and fine-tuning is essential before integrating any approach into a larger SV calling framework. In addition, we are aware that alternative strategies for phased consensus sequence generation may exist and that our implementations simplify some of the strategies implemented in previous SV callers. Our results are therefore not indicative of any existing tools’ performances, even if the implemented strategies are similar.

In conclusion, this work sheds light on several factors impacting different strategies for the generation of consensus sequences and highlights the importance of making informed choices during algorithm development. It provides a foundation for further improving our SV calling abilities and may serve as a reference for developers in the field of SV detection.

## Data Availability

We used publicly available data from the Genome in a Bottle project [Hansen et al., 2026], the human pangenome project [Liao et al., 2023] and data released by Oxford Nanopore (https://epi2me.nanoporetech.com/category/data-releases/). Download links are listed in the Supplementary Notes.

## Competing interests

No competing interest is declared.

## Author contributions statement

T.M. and B.K. conceived the study. R.L. implemented the a priori phasing and T.M. implemented the remaining strategies. T.M. conducted evaluations. All authors analysed the results and wrote and reviewed the manuscript.

## Acknowledgements

This work was in part funded by the Deutsche Forschungsge-meinschaft (DFG, German Research Foundation) – 400728090 (FOR2841); 324392634 (TRR221).

## Notes

### Competing Interest Statement

The authors have declared no competing interest.

