## Supplementary Material for "Evaluating local assembly strategies for human long-read structural variant calling"

### 1 Supplementary Notes

#### 1.1 Artificial chr1 haplotypes with random SNPs and uniformly distributed SVs

We introduced a total of 16000 SV loci in non-telomere and non-centromere regions of two chromosome 1 (chr1) copies from the T2T-CHM13 reference genome. We inserted 3200 deletions, duplications and inversion each with a heterozygous-to-homozygous ratio of 2 : 1 and additionally introduced compound heterozygous loci, i.e., loci with two variant alleles that differ from each other and the reference genome. Our compound heterozygous loci comprise either two overlapping deletions of different sizes ($n = 3200$) or an inversion on one and a deletion on the other allele in a distance of 100 bp ($n = 3200$). To further simulate single-nucleotide variants (SNVs), we randomly substituted single base positions. We chose substitution probabilities to yield an expected SNV density of one per kilobase and a targeted heterozygosity ratio [**Samuels2016**] of 1.8.

#### 1.2 Output categories

For homozygous variants, we distinguish between instances of **(A)** a single consensus sequence, **(B)** two distinct consensus sequences or **(C)** no consensus sequences being generated. In the case of heterozygous variants we distinguish between **(D)** two sequences aligning to different haplotypes, **(E)** a single sequence, **(F)** two sequences aligning to the same haplotype and **(G)** no output.

#### 1.3 Precision and Recall

In section *Performance Evaluation* we define precision and recall based on output categories only for heterozygous variants.
We make this distinction because there is an unequal number of homozygous and heterozygous loci and homozygous loci are expected to produce only a single consensus sequence, which assigns higher importance to heterozygous variants if the precision and recall were calculated across all loci.

For homozygous loci, the precision and recall are analogously defined as

$$\text{Precision} = \frac{|A| + |B|}{|A| + 2 \cdot |B|} \quad \text{and}$$
$$\text{Recall} = \frac{|A| + |B|}{|A| + |B| + |C|}.$$

#### 1.4 Phasing score for k-mer based phasing

The Ref of all selected k-mer pairs are partitioned into two haplotypes by optimizing a global phasing score $\mathcal{P} = \sum_{R_i \in \mathcal{R}} P_{R_i}$, where $\mathcal{P}_{R_i}$ is incremented by 1 for each two $k$-mer pairs $(\text{Ref}_1, \text{Alt}_1)$ and $(\text{Ref}_2, \text{Alt}_2)$ occurring in the same read $R_i$ and with the appearing alleles assigned to the same haplotype. $\mathcal{P}_{R_i}$ is decremented by 1 for each two $k$-mer pair occurrences in $R_i$ where the appearing alleles are assigned to different haplotypes. Initially, we iteratively assign the $k$-mer pairs to the haplotype that maximizes the current global phasing score $\mathcal{P}$. Subsequently, we check if the score can be increased by flipping the haplotype assignment of a single pair. While this is possible, the $k$-mer pair that yields the largest improvement is flipped. Finally, we assign the input reads to haplotypes based on the majority vote of $k$-mers they contain.

More formally defined: For a variation site $V_i \in \mathcal{V}$ a read $R_k \in \mathcal{R}$ can either support haplotype 1 ($h(V_j, R_i) = 1$), haplotype 2 ($h(V_j, R_i) = -1$) or carry no information ($h(V_j, R_i) = 0$). The global phasing score $\mathcal{P}$ is calculated as $\mathcal{P} = \sum_{V_{i,j} \in \mathcal{V}} \sum_{R_k \in \mathcal{R}} h(V_i, R_k) * h(V_j, R_k)$ for each possible combination of variation sites $i$ and $j$.

#### 1.5 Consensus sequence quality

We evaluate consensus sequence quality by aligning created sequences against the ground truth for the two haplotypes present in the genome. For real data, we assume that the SV calls in the GIAB v5.0q SV callset represent the ground truth and reconstruct the allele sequences from these calls in combination with flanking sequences extracted from the T2T-CHM13 reference genome.

To determine the quality of an assigned consensus sequence $C_i$, we count the number of matched bases $n_b$ and calculate the error rate as $1 - n_b/|C_i|$. Qualities of consensus sequences generated from real data is shown in Figure 1.

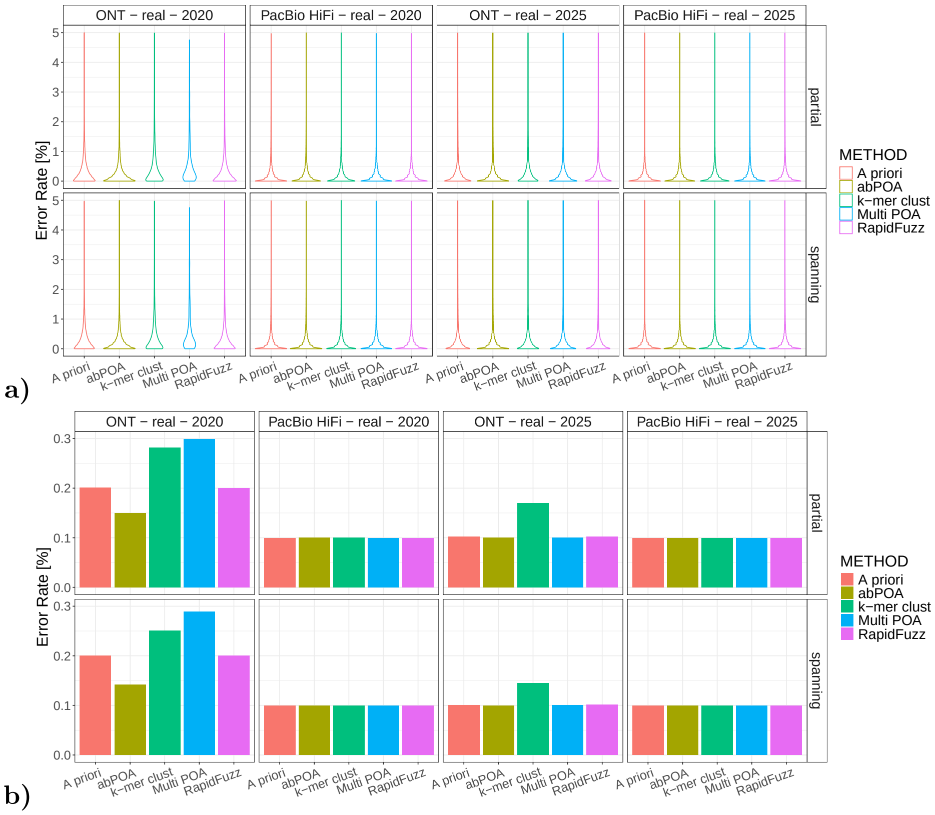

Supplementary Figure 1: Error rates of true positive consensus sequences generated with local alignment of the $\mathcal{S}^+$ input sequence set (partial) and global alignment of the $\mathcal{S}$ input sequence set on real data. Shown are the error rates for coverage 32.5×. **a)** Violin plot of all error rates per strategy. **b)** Median error rate for each strategy.

#### 1.6 K-mer size for k-mer based clustering

For the $k$-mer clustering approach described in *Similarity-based sequence clustering* we chose $k$ based on performance achieved for different values of $k$ on the artificial chr1 dataset (ONT: $k = 11$, HiFi: $k = 21$):

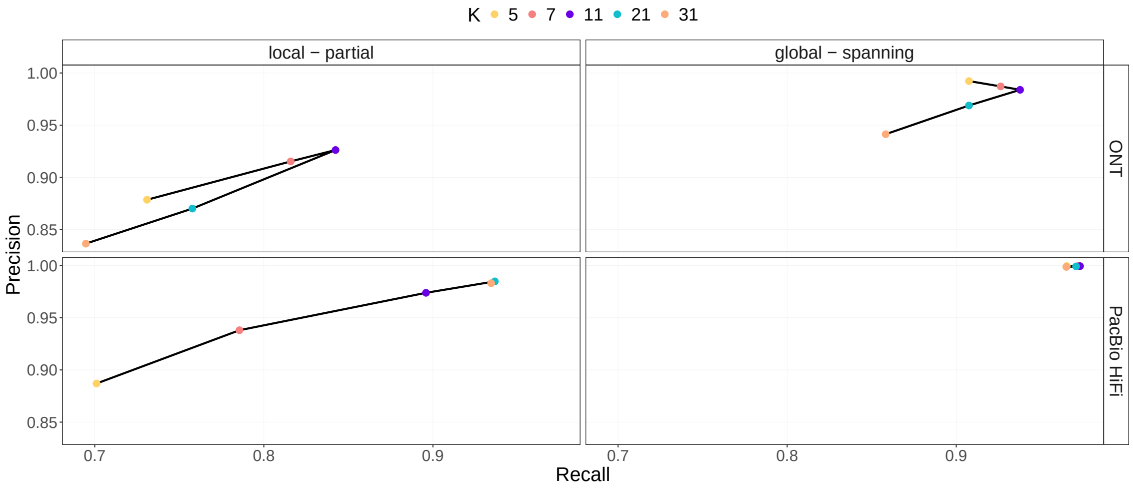

Supplementary Figure 2: Precision and recall for each tested value of $k$ in the $k$-mer based clustering approach.

We observe a similar behaviour for the error rates of generated consensus sequences compared to the ground truths.

#### 1.7 Impact of different SV contexts on consensus sequence generation

We checked all SV regions from our GIAB callset against T2T-CHM13 annotations for centromeres, telomeres, segmental duplications, tandem repeats, simple repeats and rDNA to check whether overlap with any of these repeat classes has a significant impact on the performance of any of our strategies.
If multiple different repeat types overlap with the same variant regions, we prioritize annotation as follows:

Telomere > CenSat > rDNA > Simple Repeat > Tandem Repeat > Segmental Duplication

While there is some variance in performance of the different strategies within each repeat class, the overall effect between different repeat types is much more prominent (Figures 3 and 4).

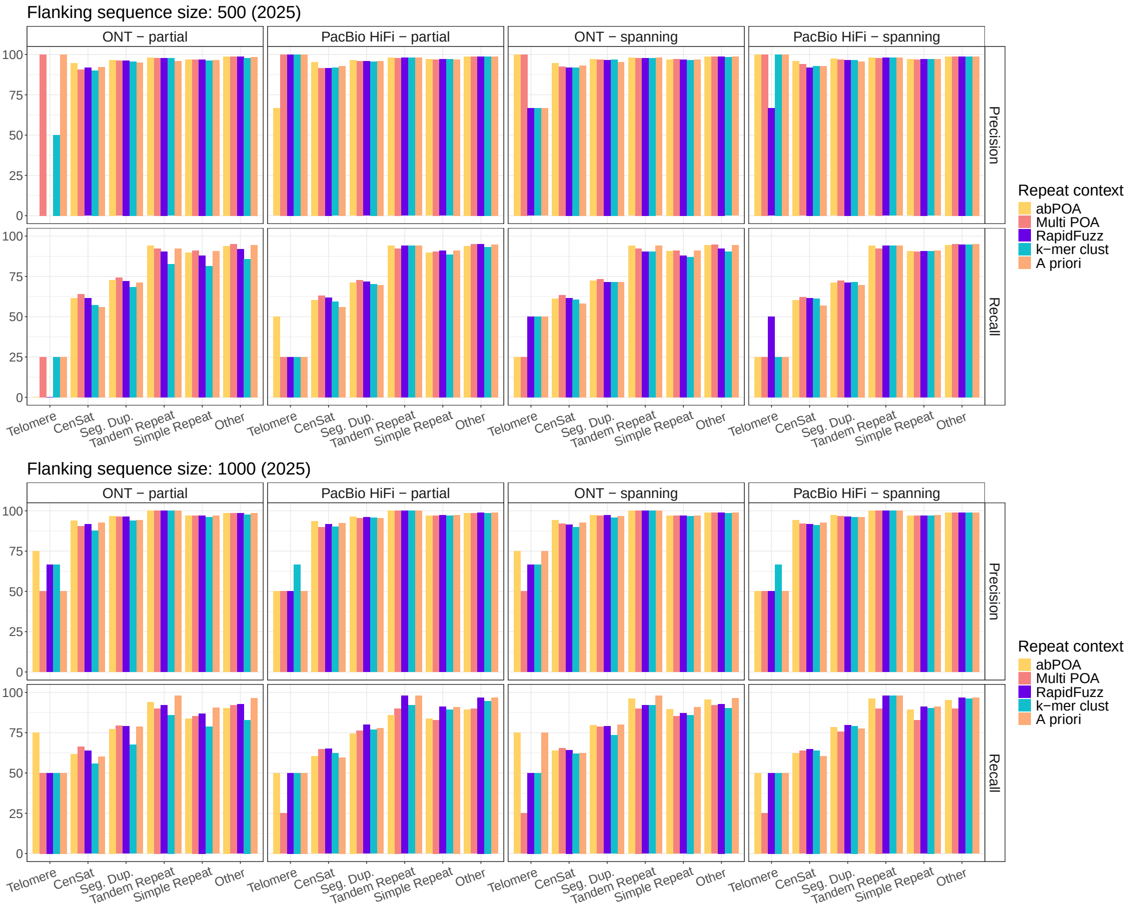

Supplementary Figure 3: Precision and recall for all methods in different repeat contexts on `ONT 2025` and `HiFi 2025` data using input sequence sets $\mathcal{S}$ (spanning) or $\mathcal{S}^+$ (partial). Coverage: 32.5×.

| | CenSat | Other | Segmental Duplication | Simple Repeat | Tandem Repeat | Telomere |
|---|---|---|---|---|---|---|
| 500 | 2719 | 6318 | 589 | 3841 | 35 | 4 |
| 1000 | 2122 | 5992 | 480 | 3692 | 34 | 4 |

Table 1: Number of variants overlapping each repeat class for different margin sizes.

Repeats seems to have little influence on the quality of the correctly generated consensus sequences (see Figure 5). Only among the four SV loci overlapping telomere regions do we observe an elevated error rate.

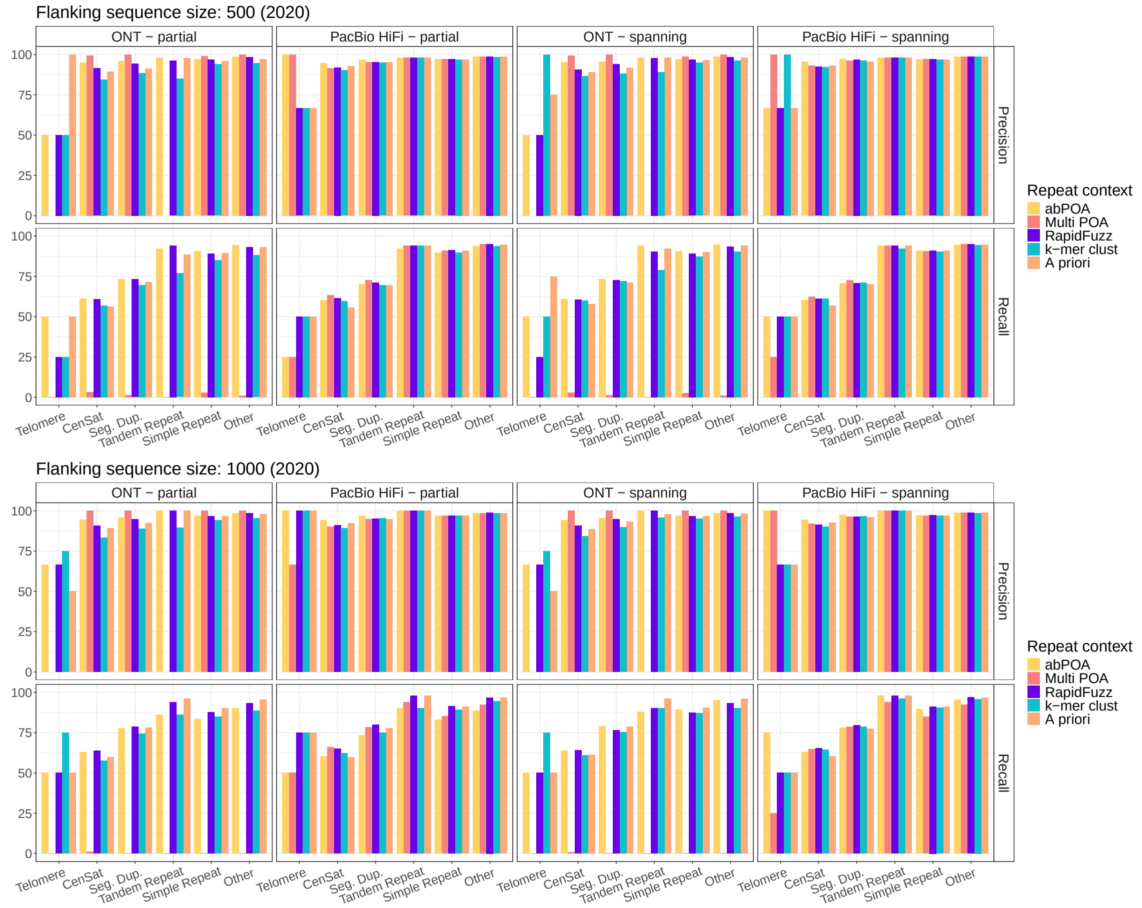

Supplementary Figure 4: Precision and recall for all methods in different repeat contexts on `ONT 2020` and `HiFi 2020` data using input sequence sets $\mathcal{S}$ (spanning) or $\mathcal{S}^+$ (partial). Coverage: 32.5×.

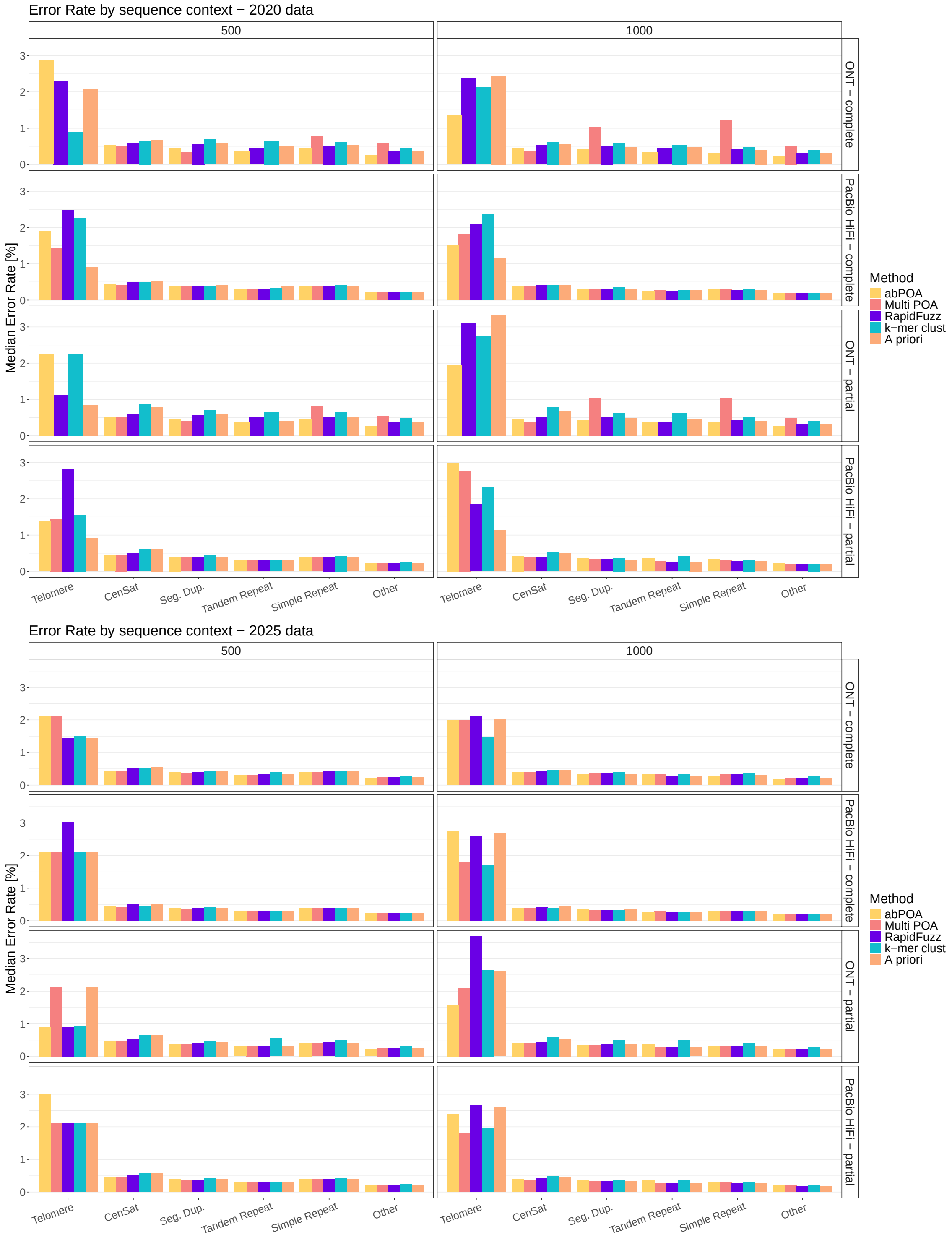

Supplementary Figure 5: Median error rate of generated consensus sequences for all methods for different repeat context.

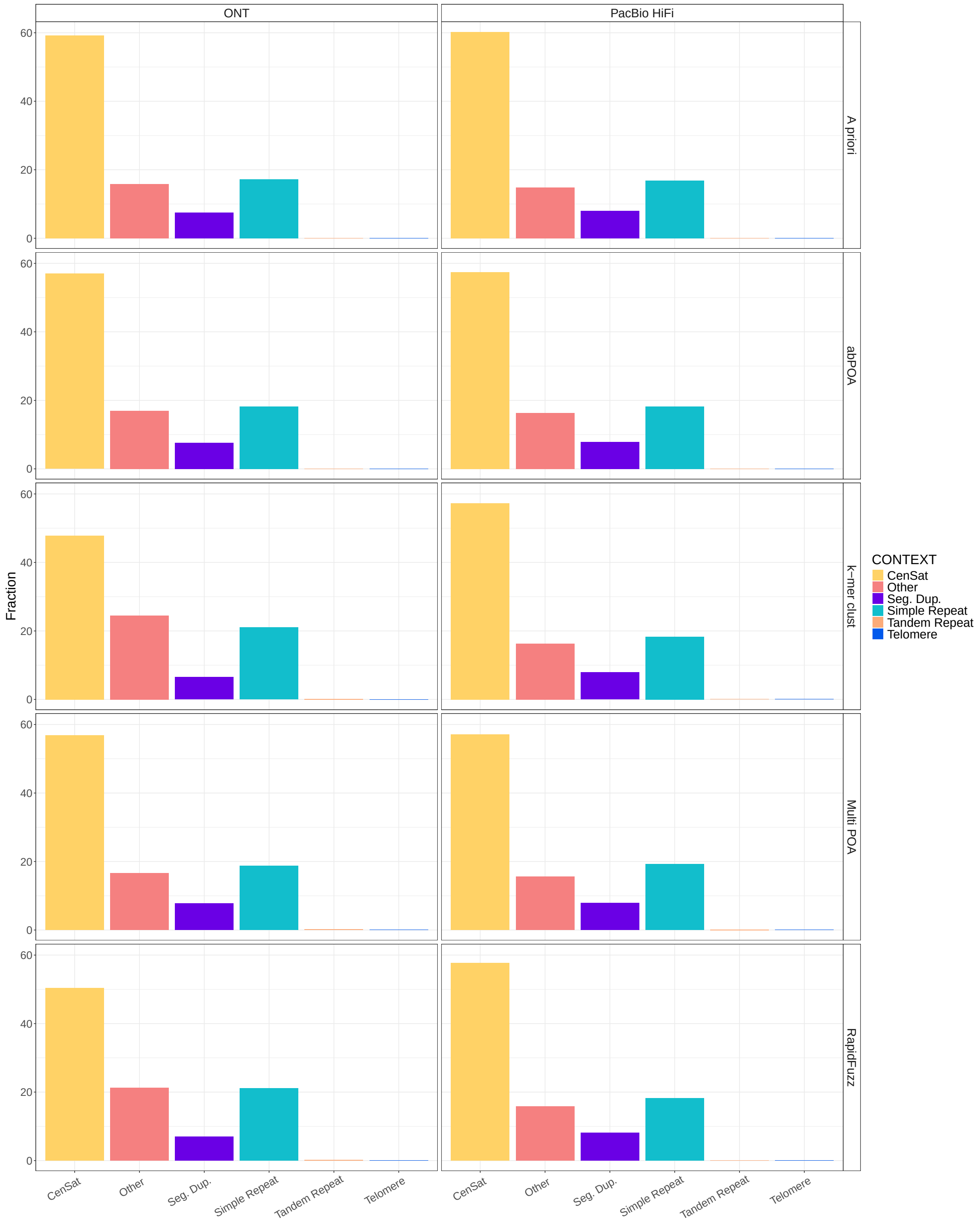

Supplementary Figure 6: Fraction of false negatives per method within different repeat contexts. `HiFi 2025` and `ONT 2025`, 32.5× coverage, input set $\mathcal{S}$ with global alignment. Columns: Sequencing technology. Rows: Different consensus strategies. For most strategies, between 50% and 60% of false negatives are located within centromere regions.

#### 1.8 Analysis of breakpoint signals in GIAB SV regions

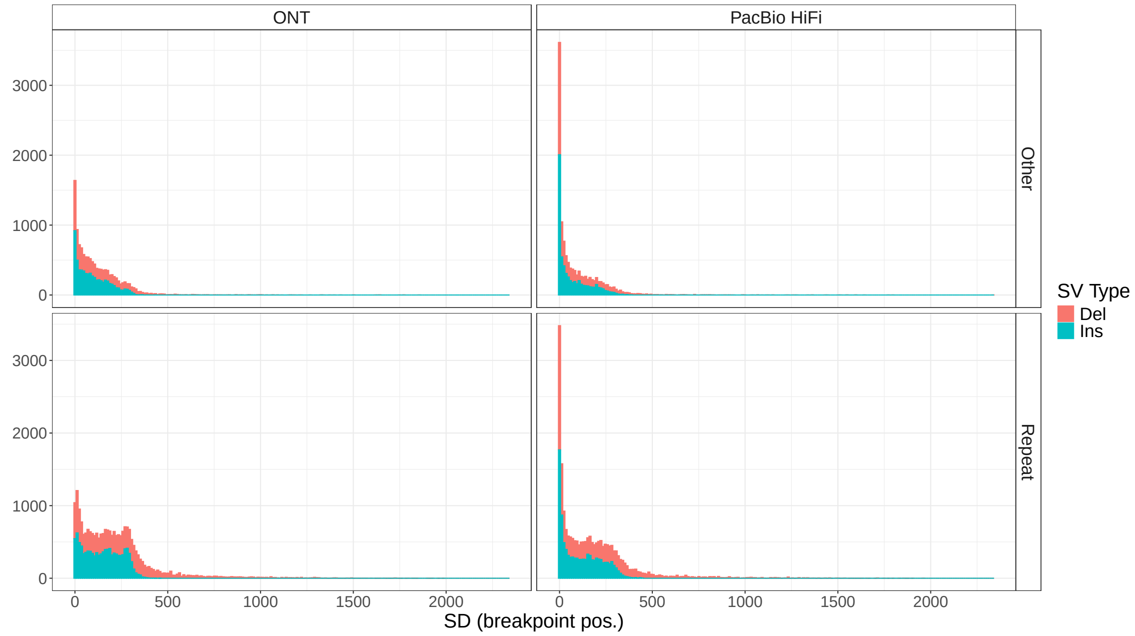

Supplementary Figure 7: Standard deviations of positions of all breakpoint positions found in an SV region, divided by SV type. Columns: Sequencing technology (2025 data). Rows: Repeat context.

We analysed the BAM files of the 32.5× real PacBio HiFi and ONT data (`ONT 2025` and `HiFi 2025`) to determine the variability in observed breakpoint positions within the filtered GIAB SV regions. We parsed alignments overlapping each GIAB SV region in a given BAM file and extracted all insertion and deletion signals contained in the CIGAR strings, as well as split or clipped alignments. We then calculate the standard deviation of all breakpoint positions in the region, as an estimate of the uncertainty of SVs called in this region (see Figure 7).

#### 1.9 Runtime analysis

We ran consensus sequence generation on `HiFi 2025` and `ONT 2025` data at 32.5× coverage using input sets $\mathcal{S}$ and $\mathcal{S}^+$ with local and global alignment, respectively. Runtime was measure for every individual consensus sequence generation and then averaged across all SV loci (see Figure 8). A priori phasing was excluded, since the workflow setup does not allow for a fair comparison of consensus sequence generation including the phasing step.

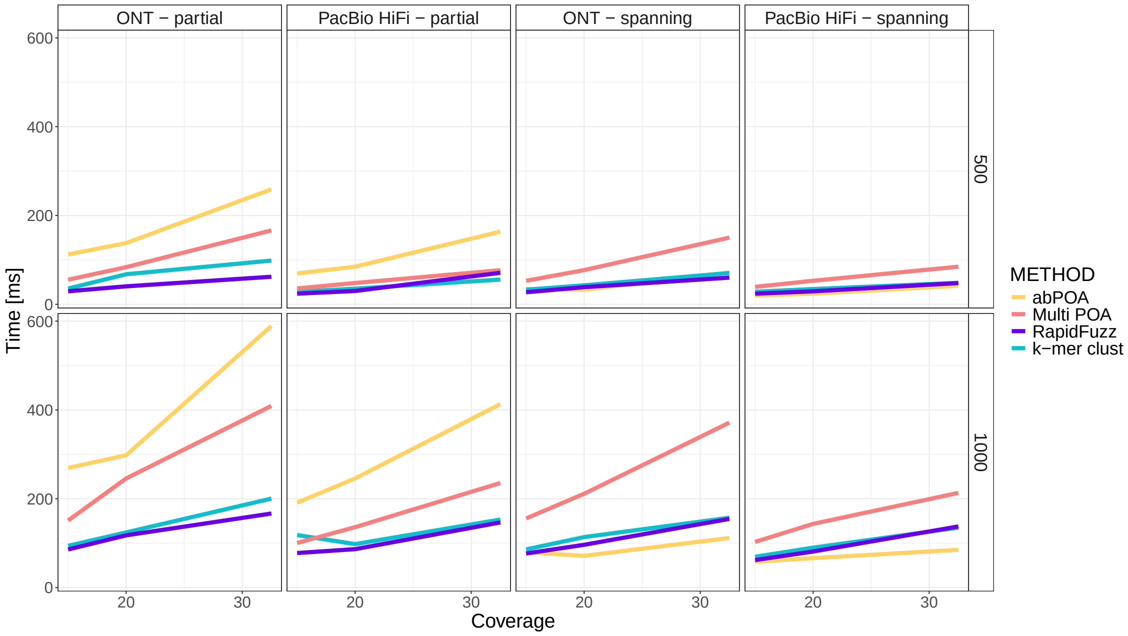

Supplementary Figure 8: Average runtime for consensus sequence generation at a single locus. Columns: Sequencing technology and input sets used. Rows: Flanking region size.

#### 1.10 Selection of alignment parameters

We ran consensus sequence generation on our uniformly simulated chromosome 1 data using different sets of alignment parameters. Based on overall performance, we decided on a edit-distance-like scoring scheme (`match: 1, mismatch: -1, gap open: -1, gap extend: -1`) for further evaluation. Precision and recall for all methods on different sequencing technologies with different input sets are shown in Figure 9. Averaged F1 scores across all methods are shown in Figure 10.

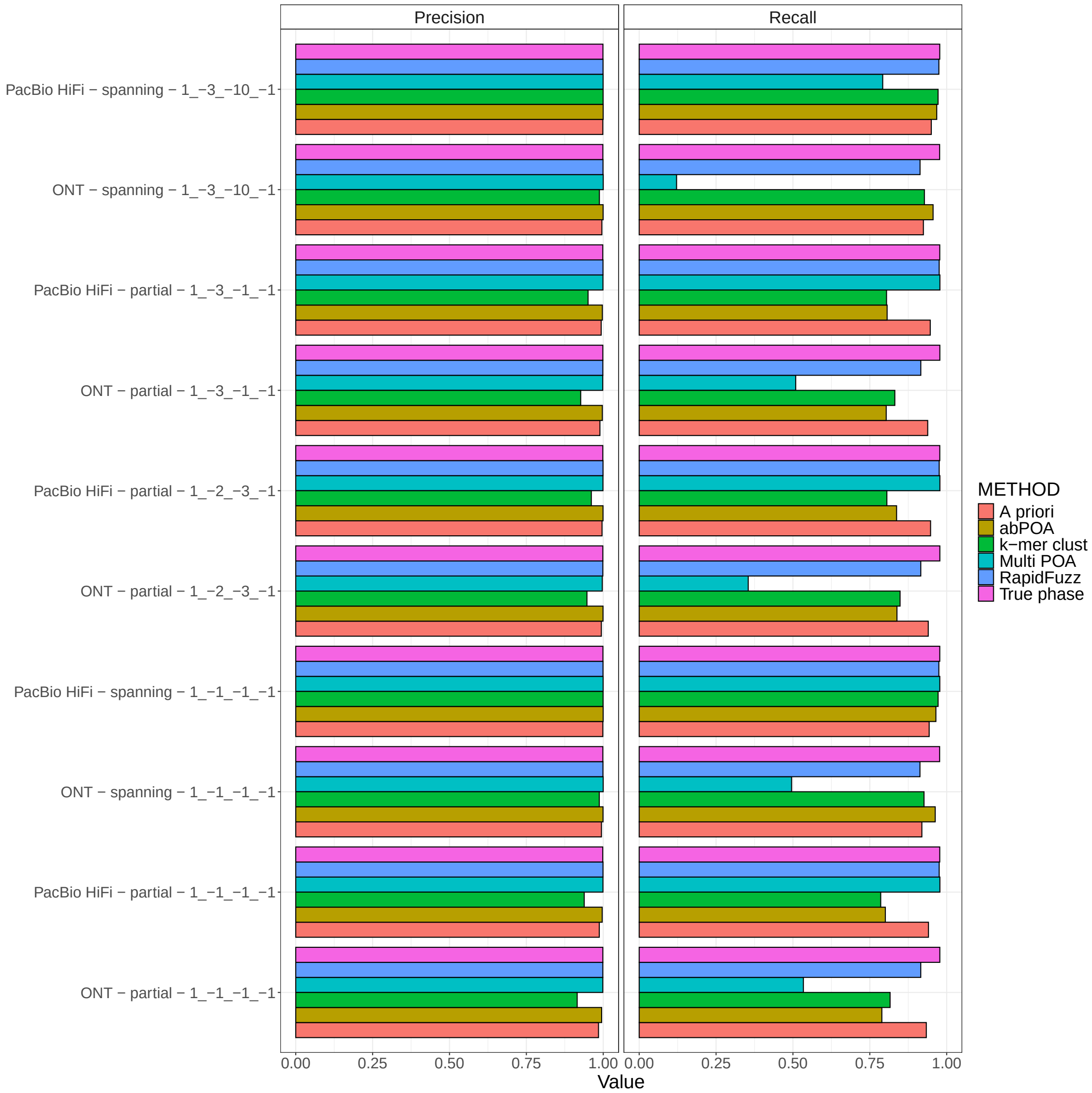

Supplementary Figure 9: Columns: Precision and Recall based on group counts. Y-axis: Sequencing technology - input read set - alignment parameters.

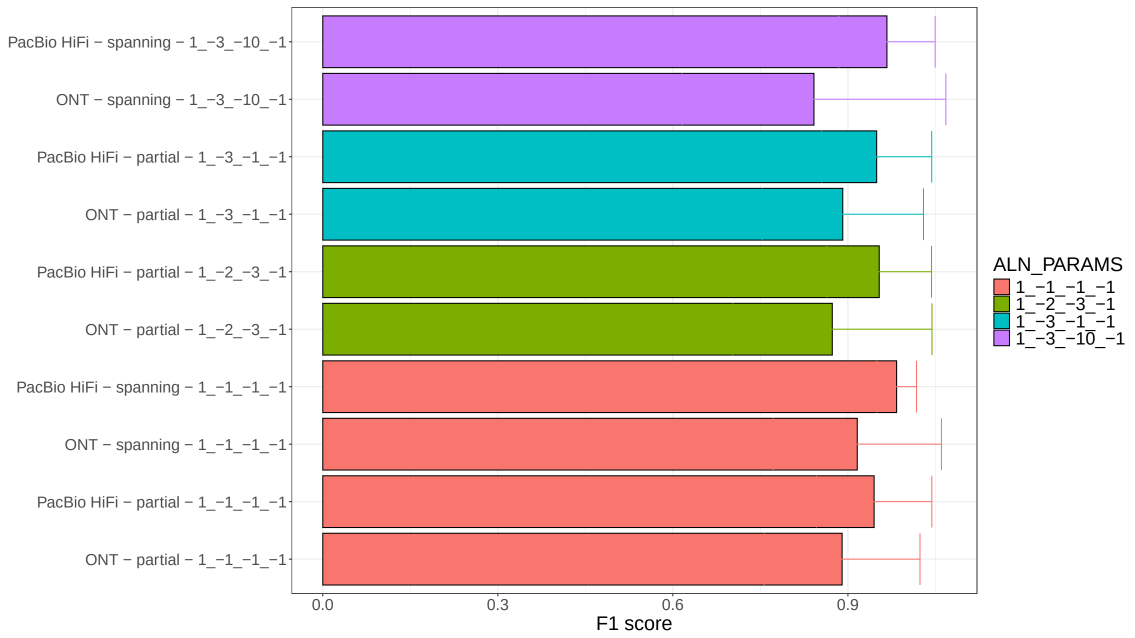

Supplementary Figure 10: Same information as in Figure 9, condensed by calculating F1 scores from precision and recall and averaging over all methods for each combination of sequencing technology, input set and alignment parameters.

#### 1.11 Data links

##### VCF files

VCF files containing HG002 variant calls from GIAB:

- SVs: https://42basepairs.com/download/web/giab/release/AshkenazimTrio/HG002_NA24385_son/v5.0q/HG002_CHM13v2.0_v5.0q_stvar.vcf.gz
- small variants: https://42basepairs.com/download/web/giab/release/AshkenazimTrio/HG002_NA24385_son/v5.0q/HG002_CHM13v2.0_v5.0q_smvar.vcf.gz

Recent sequencing data for HG002:

- ONT 2025: s3://ont-open-data/giab_2025.01/basecalling/sup/HG002/
- HiFi 2025: https://s3-us-west-2.amazonaws.com/human-pangenomics/T2T/scratch/HG002/sequencing/hifirevio/m84011_220902_175841_s1.hifi_reads.fastq.gz

Older ONT sequencing data:

- s3://ont-open-data/gm24385_2020.11/analysis/r9.4.1/20201026_1644_2-E5-H5_PAG07162_d7f262d5/guppy_v4.0.11_r9.4.1_hac_prom/basecalls.fastq.gz

Older HiFi data:

- https://s3-us-west-2.amazonaws.com/human-pangenomics/NHGRI_UCSC_panel/HG002/hpp_HG002_NA24385_son_v1/PacBio_HiFi/15kb/m64012_190920_173625.Q20.fastq
- https://s3-us-west-2.amazonaws.com/human-pangenomics/NHGRI_UCSC_panel/HG002/hpp_HG002_NA24385_son_v1/PacBio_HiFi/15kb/m64012_190921_234837.Q20.fastq
- https://s3-us-west-2.amazonaws.com/human-pangenomics/NHGRI_UCSC_panel/HG002/hpp_HG002_NA24385_son_v1/PacBio_HiFi/20kb/m64011_190830_220126.Q20.fastq
- https://s3-us-west-2.amazonaws.com/human-pangenomics/NHGRI_UCSC_panel/HG002/hpp_HG002_NA24385_son_v1/PacBio_HiFi/20kb/m64011_190901_095311.Q20.fastq

### 2 Supplementary Figures

#### 2.1 Supplement to Figure 2b (main text)

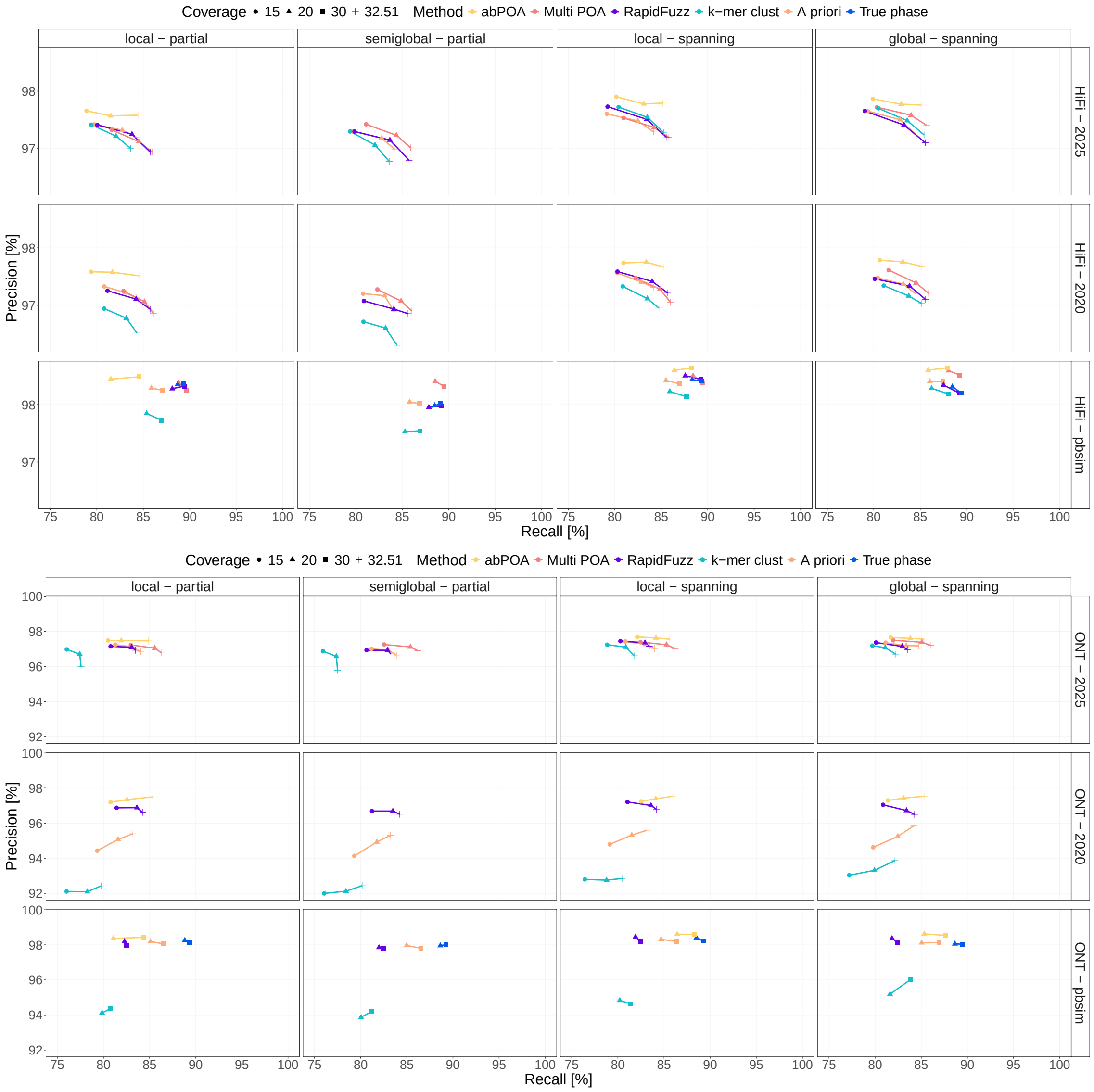

Supplementary Figure 11: Precision-Recall curves, including `HiFi 2025` and `HiFi - pbsim`. Real data was subsampled to 32.5× (coverage of full `HiFi 2025` BAM file).

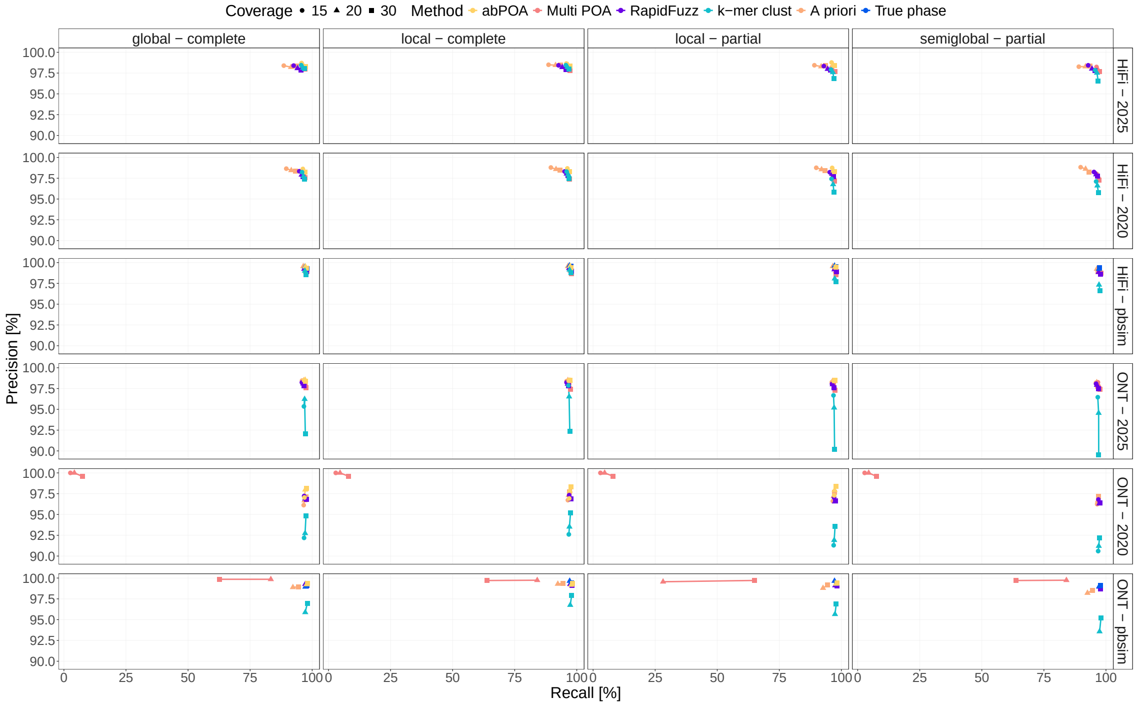

Supplementary Figure 12: Precision-recall curves only for homozygous SV loci.

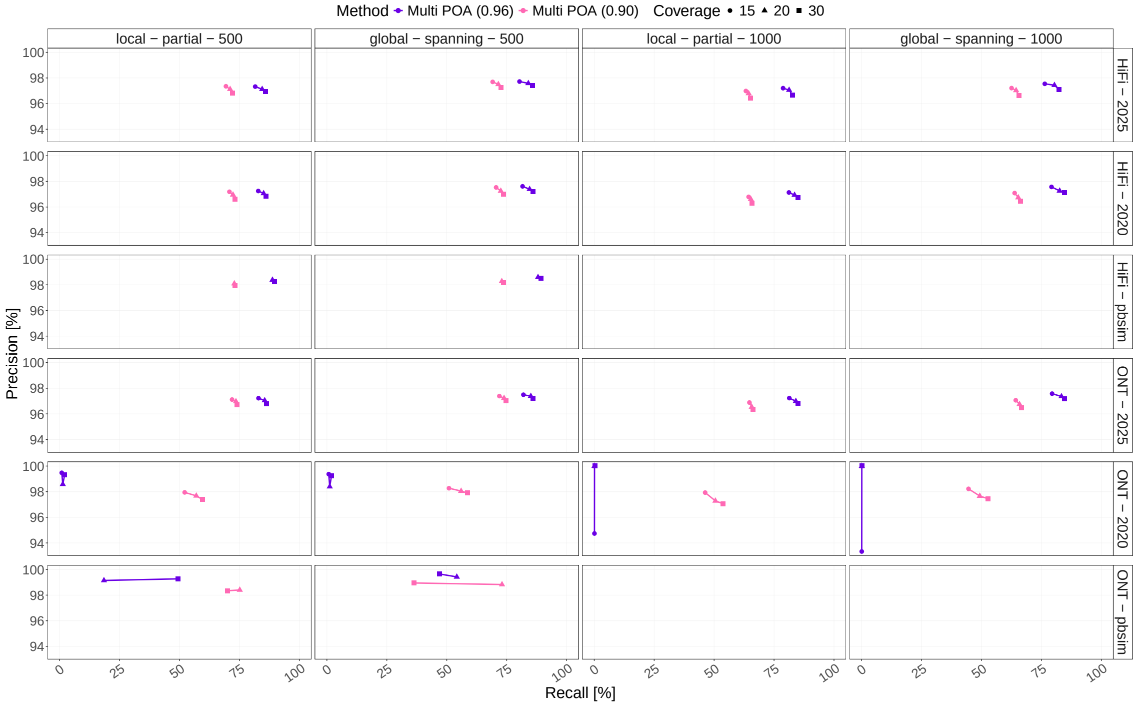

Supplementary Figure 13: Precision-recall curves for Multiple POA strategy with two different thresholds – 0.96 (default) and 0.90 – for internal read-to-graph alignment.

#### 2.2 Supplement to Figure 3 (main text)

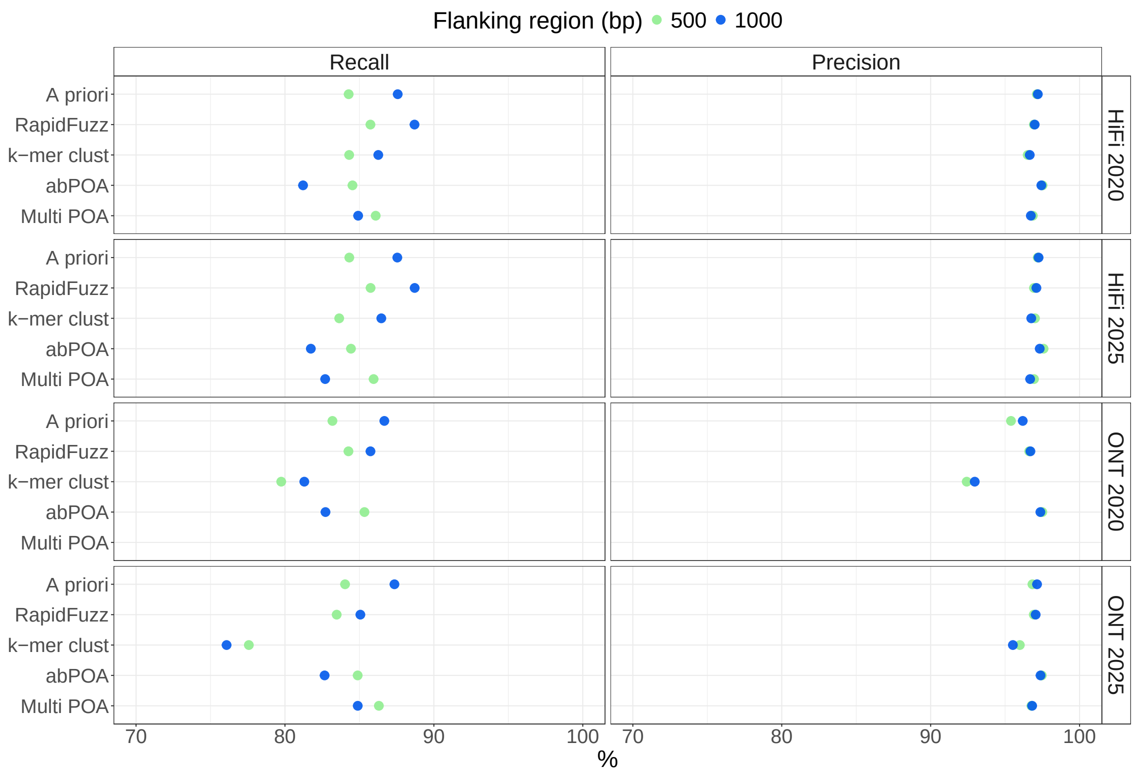

Supplementary Figure 14: Figure 3a with input sequence set $\mathcal{S}^+$ and local alignment.

#### 2.3 Flanking sequence size impact on simulated data

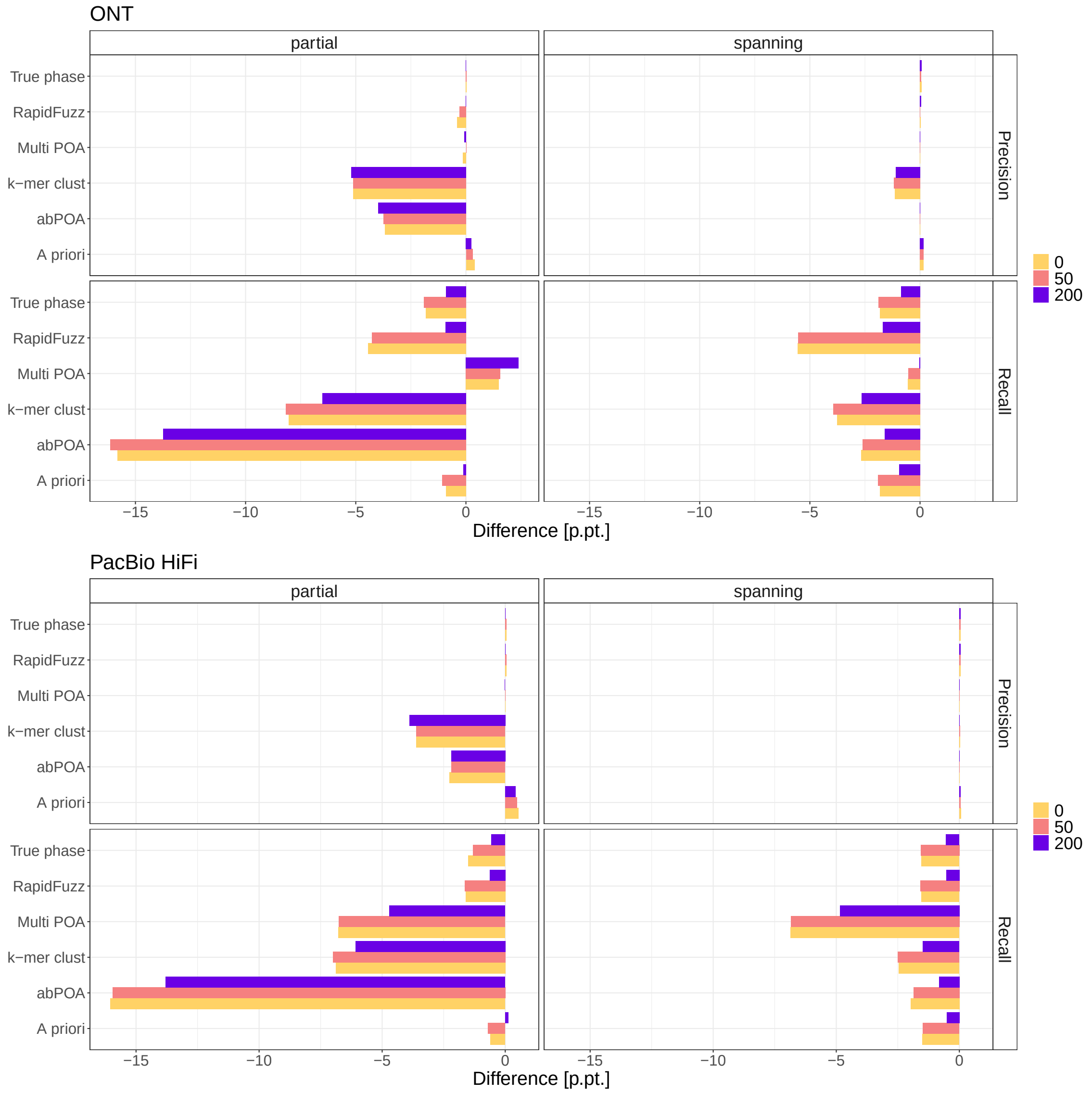

Supplementary Figure 15: Change of precision and recall in percentage points when increasing flanking region size $f$ from 500 to 1000 on the uniformly simulated chromosome 1 data set. Simulated with different degrees of uncertainty, adding Gaussian noise with an SD of 0, 50, or 200 to the start and end positions of the regions placed around breakpoints. Contrary to observations on real data, recall decreases slightly with increasing flanking region size. Possibly due to longer sequences containing more noise, negatively impacting similarity calculations. The effect is diminished by increased simulated breakpoint uncertainty (200), suggesting that the positive recall effect observed on real data may be due to breakpoint uncertainty in the callset, or differences in SV placement between read-alignment and assembly-based methods.

#### 2.4 F1-score by simulated SV type

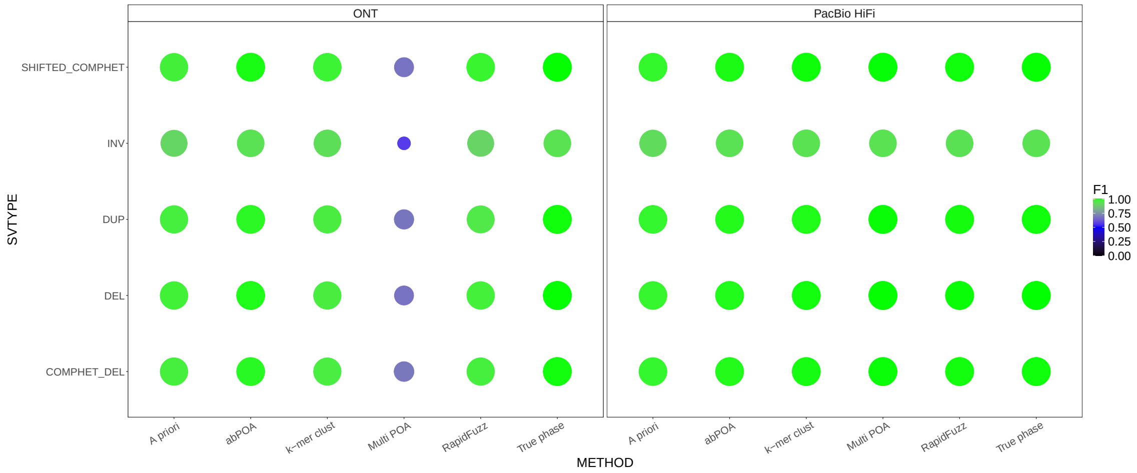

Supplementary Figure 16: F1 score per method for each variant type. Smaller point size indicates lower F1 score. Del: deletion, Inv: inversion, Dup: duplication, Comphet_Del: compound heterozygous deletion, Shifted_Comphet: non-overlapping heterozygous deletion-inversion locus. Input read set: $\mathcal{S}$, global alignment, 30× coverage, alignment parameters `1, -1, -1, -1`.

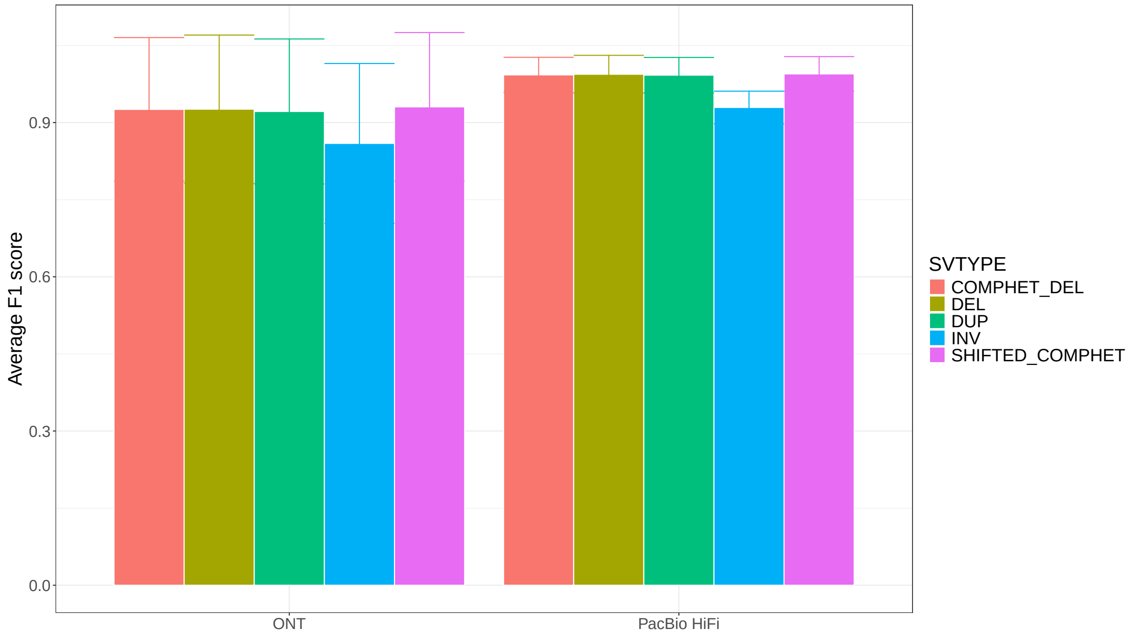

Supplementary Figure 17: Same information as in Figure 16, condensed by averaging across methods. Inversions are slightly harder to resolve accurately.
